# Sulfide is a keystone metabolite for gut homeostasis and immunity

**DOI:** 10.1101/2025.03.06.641928

**Authors:** Victor I. Band, Inta Gribonika, Apollo Stacy, Nicolas Bouladoux, Shreni Mistry, Andrew Burns, P. Juliana Perez-Chaparro, Joanna Chau, Michel Enamorado, Motoyoshi Nagai, Vanya Bhushan, Dominic P. Golec, Pamela L. Schwartzberg, Suchitra K. Hourigan, Aleksandra Nita-Lazar, Yasmine Belkaid

## Abstract

Hydrogen sulfide is a gaseous, reactive molecule specifically enriched in the gastrointestinal tract. Here, we uncover a non-redundant role for sulfide in the control of both microbial and immune homeostasis of the gut. Notably, depletion of sulfide via both pharmaceutical and dietary interventions led to a profound collapse of CD4 T cells in the ileum of the small intestine lamina propria and significant impact on microbial ecology. As a result, mice with reduced sulfide within the gut were deficient in their ability to mount T cell dependent antibody responses to oral vaccine. Mechanistically, our results support the idea that sulfide could act directly on CD4 T cells via enhanced AP-1 activation, leading to heightened proliferation and cytokine production. This study uncovers sulfides as keystone components in gut ecology and provides mechanistic insight between diet, gut sulfide production and mucosal immunity.

## Main Text

Sulfur compounds acted as the earliest electron acceptors, and are highly reactive, able to modify a range of organic compounds (*1, 2*). Hydrogen sulfide is proposed to have played an essential role as a redox molecule for early life, thriving in an oxygen-free atmosphere abundant in sulfur gases (*3, 4*). The gradual oxygenation of the atmosphere resulted in elimination of this sulfide pool (*2*). Today, the primitive metabolic processes involved in sulfur metabolism remain observable in extreme anaerobic environments such as in prokaryotes living in geothermal hot springs (*5*). Very low concentrations of hydrogen sulfide are also produced in mammals and other higher organisms and have been recently shown to play a key role in various biological processes (*6*) including memory formation, smooth muscle contraction and protection against oxidative stress (*7-9*). In these contexts, the action of sulfide depends on nano-molar to low micro-molar levels of sulfide production (*10*), while higher concentrations of sulfide are known to be toxic (*11*).

A notable exception is the gastrointestinal (GI) tract, where very high sulfide concentrations have been reported reaching up to 3 mM (*12*). The uniquely enriched sulfide levels within the gut result from several factors, notably the combination of an anaerobic environment with high abundance of prokaryotic members of the microbiome. While the microbiome represents a significant source of sulfide, nearly half of gut sulfide originates from the host epithelium (*13*). Dietary factors are major determinants of gut sulfide production, as protein intake can drive fecal sulfide production (*14*), specifically due to sulfide production from sulfur containing amino acids (SAAs). As such, sulfide level can be greatly impacted in low protein diet such as in vegetarians/vegans and more particularly in the context of malnutrition (*14, 15*). In humans, sulfide levels can also be significantly impacted by commonly used drugs. For instance, bismuth subsalicylate (BSS), an over-the-counter anti-diarrheal drug, is known to deplete gut sulfides (*16*). These exposures result in sulfide levels within the gut that vary greatly across individuals and over time based on diet, microbiome composition and drug exposure (*14, 17*).

The GI tract is also the primary site of exogenous exposures ranging from microbes, both symbiotic and pathogenic, to dietary antigens. To ensure regulated responses to these complex demands, the gut is equipped with a specialized immune system, with over half of the total immune cells of the host residing within this compartment (*18*). The extreme levels of sulfide found within the gut raise fundamental questions: How do immune cells adapt to and potentially benefit from high sulfide concentrations? What roles do sulfides play in mucosal, microbial, and immune homeostasis? How do fluctuations in sulfide metabolism, influenced by diet or drug consumption, impact mucosal immunity?

Here, we explore the role of sulfides in gut immune and microbial homeostasis. Surprisingly, our results reveal that at the sites of highest sulfide levels, such as the ileum, T cells not only withstand these conditions but, as we demonstrate, actively rely on this metabolite to sustain their survival and function. Collectively, we reveal a non-redundant role for sulfides in gut immunity and microbial ecology, providing important insights into the collateral effects of commonly used drugs and a potential explanation for the collapse of mucosal immunity and response to oral vaccines in undernourished populations.

## Results

### Depletion of gut sulfides selectively reduces CD4 T cells in the ileum

While previous reports documented high level of sulfides within the gut, characterization of sulfide levels within defined gut compartment had not been assessed. Testing of free sulfide concentrations across the small intestinal lumen revealed a gradual concentration increase, reaching maximal levels in the distal ileum with over 5000μM luminal sulfide (**Fig 1B**). This indicates that the highest levels of sulfide are found in the portion of the small intestine associated with the greatest microbial load and Peyer’s patch (PP) size and density (*19*). This observation pointed to a potential role for this gas in immune and microbial ecology.

**Figure 1.**
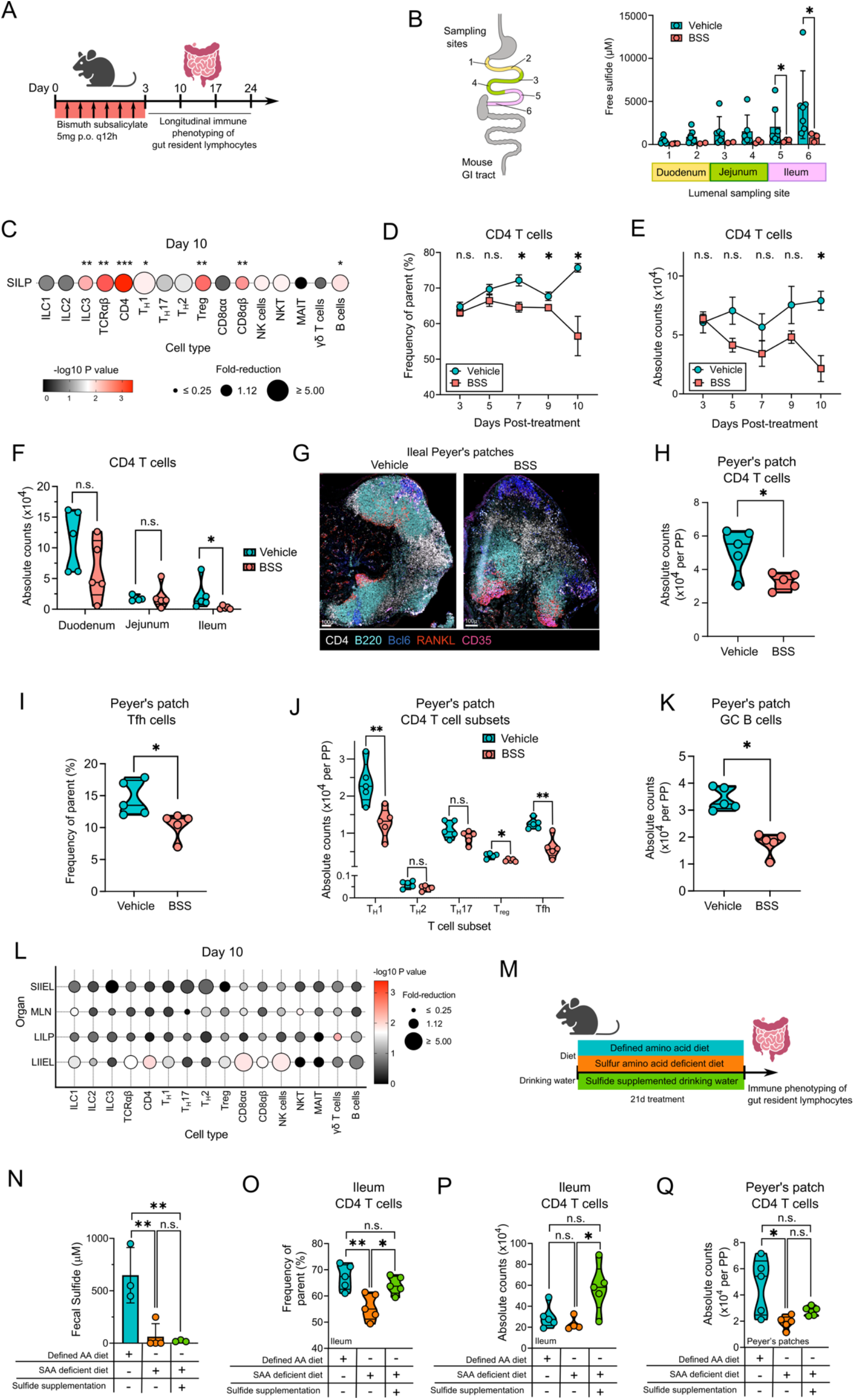
Gut CD4 T cells are reduced after depletion of gut sulfides using bismuth or diet. (**A**) Schematic of bismuth treatment protocol. Mice were treated with 5mg BSS every 12 hours for 72 hours, then assessed at day 10 for immune phenotyping. (**B**) Measurements of free sulfide at various locations in the small intestine, as noted on the left. Right, free sulfide concentrations as measured by microsensor in the small intestinal lumen in vehicle or BSS treated mice, at 4 hours post-treatment. (**C**) Cell counts for various immune cells in the small intestine lamina propria (SILP) at day 10. Size of circles indicates the fold reduction in cells of BSS treated mice compared to vehicle and color indicates p value (n=5). (**D, E**) Frequency (D) and counts (E) of CD4 T cells from days 3 to 10 post vehicle or BSS treatment (n=5). (**F**) CD4 T cell counts in the duodenum, jejunum and ileum lamina propria of the small intestine. (**G**) Ileal Peyer’s patches of vehicle or BSS treated mice at 10 days post treatment, stained with CD4 (white) B220 (teal) Bcl6 (blue) RANKL (red) and CD35 (pink). Scale bar indicates 100μm. (**H-K**) Total counts in ileal Peyer’s patches of CD4 T cells (H), Tfh frequency (I) CD4 T cell subsets (J) or germinal center B cells (K) at day 10 of Vehicle or BSS treatment. (**L**) Cell counts for various immune cells in the small intestine intraepithelial lymphocytes (SIIEL), mesenteric lymph node (MLN), large intestine lamina propria (LILP) and intraepithelial lymphocytes (LIIEL). Size of circles indicates the fold reduction in cells of BSS treated mice compared to vehicle (n=5). (**M**) Schematic of SAA diet treatment protocol, mice were placed on a defined amino acid diet (teal), SAA deficient diet (orange) or SAA deficient diet plus NaHS in drinking water, for 21 days before quantification of immune cells in gut. (**N**) Fecal sulfide concentrations after 21 days on dietary interventions in (M). (**O**,**P)** Frequency (O) and total counts (P) of CD4 T cells in ileum lamina propria. (**Q)** Total counts of CD4 T cells per Peyer’s patch. n.s. not significant, * p < 0.05, ** p < 0.01, *** p < 0.001 as calculated by two-tailed t-test, with Benjamini-Hochberg correction for multiple comparisons (C, L).

To assess the potential role of gut sulfides in mucosal immunity, we developed a protocol to deplete gut sulfides using oral administration of bismuth subsalicylate (**Fig 1A)**. BSS dissociates into free bismuth ions in the gut, which are not absorbed but instead react with free sulfide to form the stable and non-reactive bismuth sulfide, thus reducing gut sulfide concentrations (*16*). We employed a dosing regimen (5 mg dose every 12h) meant to reproduce human dosing of BSS (*20*). Following 3 days of treatment, gut sulfide concentrations were significantly reduced in the ileum, the site of maximal sulfide concentration (**Fig 1B**) and in the feces (**Supplemental Figure 1A)**. Using this approach, we assessed the consequence of reduced sulfide levels in immune homeostasis, and more specifically on the lymphoid compartment (**Fig 1C, Supplemental Figure 1B-D**). Among all organs tested, the lamina propria compartment of the small intestine was the most significantly impacted (**Fig 1C, Supplemental Figure 1B-D**).

Notably, by 10 days post-treatment, the absolute number of small intestine CD4 T cells, CD8αβ T cells and ILC3s were significantly reduced compared to controls with minor impact on other lymphoid cells (**Fig 1C**). Among affected populations, CD4 T cells and more specifically T_H_1 cells were the most impacted as early as 7 days post treatment reaching maximum decrease by day 10 (**Fig 1D, 1E)**. In agreement with the privileged sulfide environment within the ileum (**Fig 1B**), CD4 T cells were selectively eliminated from the ileum, with ileal CD4 T cell counts reduced by >90% compared to control (**Fig 1F**).

The distal ileum represents a unique compartment of the small intestine with the highest concentration of Peyer’s patches (*21*), sites of privileged antigen sampling and germinal center (GC) responses. A key subset of CD4 T cells that drive the GC response in the Peyer’s patch are the T follicular helper cells (Tfh), which express Bcl6 and provide key signals to B cells (*22*).Within ileal Peyer’s patches, sulfide depletion impacted CD4 T cells and was associated with reduced compactness of B cell follicles and interfollicular T cell zones (**Fig 1G and 1H)**. The frequency of Tfh cells was significantly reduced in Peyer’s patches of BSS treated mice with overall numbers of Tfh cells per Peyer’s patch reduced by more than half (**Fig 1I, J)**. Further, we observed a significant decrease in GC B cells as well as T_H_1 and T_reg_ with no impact on T_H_17 or T_H_2 cells (**Fig 1J, 1K)**. On the other hand, few significant changes were observed in the large intestine lamina propria, the intraepithelial lymphocytes, or in the mesenteric lymph nodes **(Fig 1L)**. Thus, the impact of sulfide depletion is highly localized to the small intestine lamina propria, and specifically within the distal region of the ileum.

To confirm that the impact on CD4 T cells was driven by sulfide, we next employed complementary approaches to modulate sulfide levels. First, we used an alternate bismuth formulation, bismuth subnitrate, that did not contain salicylate that is a component of BSS. Using this alternative approach, we observed reduction in the absolute number of CD4 T cells within the ileum post-treatment (**Supplemental Figure 1E, Supplemental Figure 1F)**. In the gut, H_2_S is mainly produced by both the host and the microbiome from sulfide precursors such as SAAs cysteine and methionine (*23*). Thus, and to reproduce physiological settings associated with low protein intake, mice were given a diet with amino acids in place of protein, with either complete amino acids (approximating normal dietary intake) or containing amino acids with low methionine and no cysteine (SAA diet) (**Table S1, Fig 1M**). Mice on a reduced SAA diet for 2 weeks had greatly diminished free sulfides in their feces and echoing our results with BSS treatment, had reduced frequency and absolute numbers of CD4 T cells in the ileum and Peyer’s patches (**Fig 1N-Q**).

Supplementation of SAA diet by adding NaHS to drinking water did not restore fecal levels of free sulfides (**Fig 1N**), (a known limitation of NaHS supplementation (*24*)), but was sufficient to partially restore the frequency and number of CD4 T cells in the ileum and Peyer’s patches (**Fig 1O-Q**). Thus, using several independent models of gut sulfide depletion, our results uncover sulfides as a key determinant of small intestine CD4 T cell homeostasis.

### CD4 T cell collapse occurs independently of BSS-induced gut microbiome changes

The microbiome plays an important role in shaping gut immunity (*25*). To assess if sulfide control of CD4 T cells was microbiome dependent, we first tested the impact of BSS treatment on the microbiome (**Fig 2A**). While BSS treatment had no impact on alpha diversity in the feces and small intestine lumen, sulfide reduction significantly impacted fecal microbiome beta diversity as early as 3 days post-treatment (**Supplemental Figure 2A, 2B**) (**Fig 2B**). Significant differences in the microbiome beta diversity were also found at day 3 in the lumen and mucosa of the small intestine (**Fig 2C**). Several important taxa were found to be driving this compositional change throughout the gut microbiome. In the small intestine we noted a significant decrease in segmented filamentous bacteria (SFB, Candidatus *Arthromitus*) and *Muribaculum* and increase in *Akkermansia*, and within the feces a decrease in *Lactobacillus* and *Turicibacter* (**Fig 2D, Supplemental Figure 2C-E**). Of note both SFB and *Lactobacillus* have been shown to contribute to immune activation within the gastrointestinal tract (*26, 27*). We noted a profound increase in the facultative anaerobe *Enterococcus*, which has previously been observed to bloom after sulfide depletion (*28*). Metagenomic analysis revealed significant functional alterations in the microbiome after BSS treatment, including a change in glucarate and galactarate degradation and synthesis pathways which have been associated with pathogen metabolism (**Supplemental Figure 2F**) (*29*). Alteration in microbial composition following transient depletion of gut sulfide gradually decreased over time (**Fig 2B**), but significant differences persisted up to day 30 post-treatment (**Supplemental Figure 2G**). These data indicate that the sulfide depleted microbiome is significantly altered by both composition and functional capacity, which could have functional consequences.

**Figure 2.**
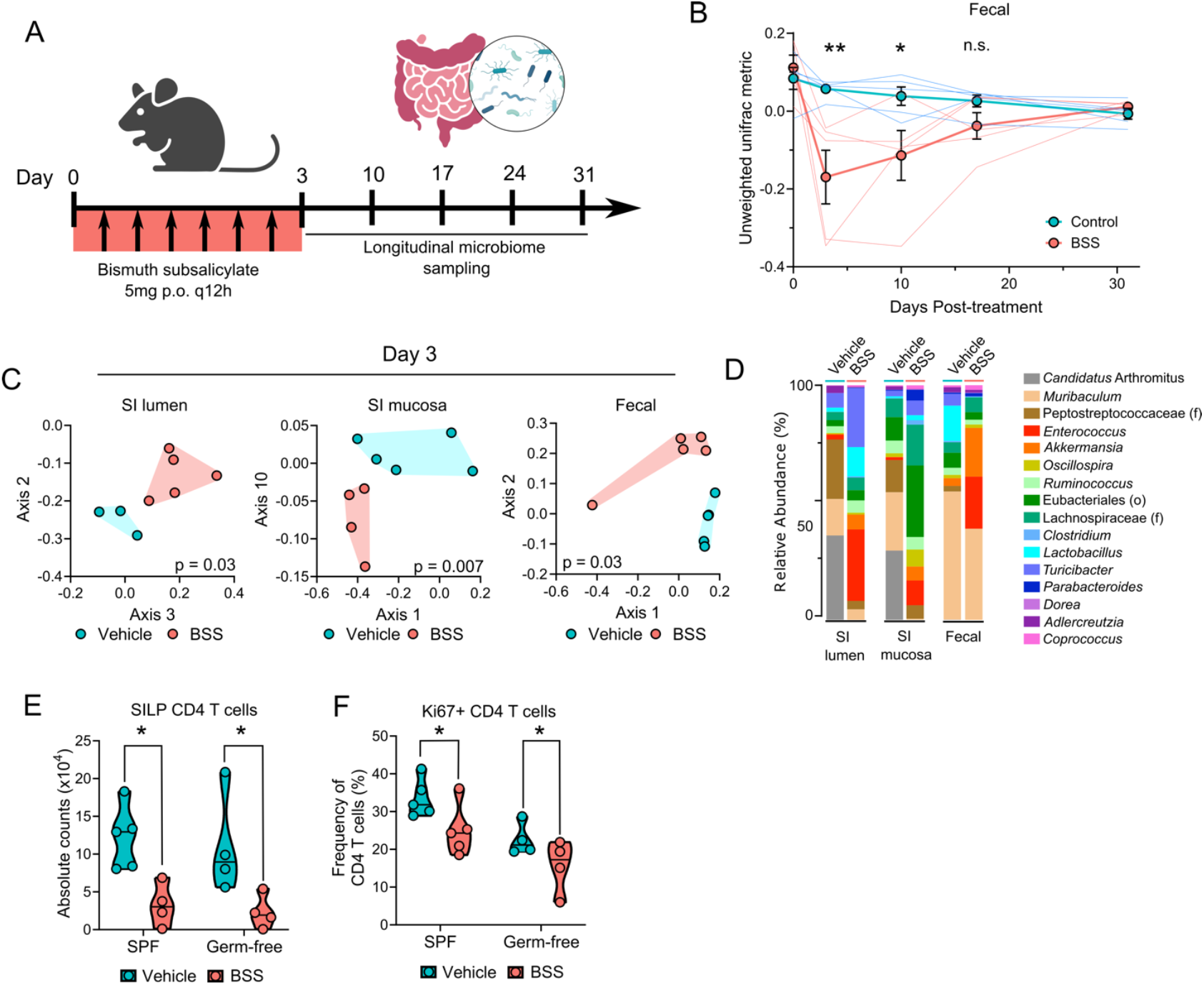
Sulfide depletion modulates gut microbiome composition and function, which does not drive CD4 T cell loss. (**A**) Schematic of microbiome sampling, mice were treated with BSS as in Fig. 1A, then microbiome compartments were sampled weekly for longitudinal samples. (**B**) Longitudinal tracking of fecal microbiome beta diversity in vehicle and BSS treated mice, as measured by Axis 2 of PCA of unweighted unifrac distance from 16S rDNA sequencing. (**C**) Microbiome sampling at day 3 post treatment of small intestinal lumen, mucosa, and feces, as measured by PCA plots of unweighted unifrac beta diversity with indicated axes. Adjusted p values calculated by PERMANOVA. (**D**) Relative abundance of top 16 identified taxa at the genus (or order/family) level in each gut compartment. (**E, F**) Flow cytometry of SILP in conventional (SPF) and germ free mice treated with vehicle or BSS, with the total numbers of CD4 T cells (E) and percent Ki67+ CD4 T cells (F) shown. n.s. not significant, * p < 0.05, ** p < 0.01, as measured by two-tailed test (E,F) or Mann-Whitney (B).

To assess whether BSS-induced microbial alteration contributed to the collapse in gut CD4 T cells, we depleted gut sulfides in mice devoid of live bacteria (germ-free). Depletion of CD4 T cells post-treatment was still detectable in germ free mice, supporting that sulfide levels can directly impact CD4 T cell homeostasis independently of its impact on the microbiota (**Fig 2E, 2F**). This prompted us to further explore the mechanism underlying the connection between high sulfide levels and CD4 T cell homeostasis and function.

### Sulfide promotes mucosal CD4 T cell activation and proliferation

To begin to address the mechanism by which sulfide impacted CD4 T cells within the gut we further characterized the phenotypic changes in T-bet^+^ T_H_1 cells, the subset that is the most impacted by gut sulfide depletion. Upon screening of a range of phenotypic markers (**Supplemental Figure 3A**), we observed a profound loss in Ki67 expression (**Fig 3A, 3B**), a marker of cell proliferation, in T-bet expressing CD4 T cells, resulting in dramatically reduced number and frequency of proliferating T-bet cells (**Fig 3C, 3D**). Intensity of T-bet expression by T_H_1 cells was also significantly reduced (**Fig 3E**). Decreased T-bet expression has been correlated with reduced ability of CD4 T cells to produce interferon gamma, which we observed was also significantly impaired after BSS treatment (**Fig 3F**). While impact in absolute number was most pronounced for T_H_1 cells, other subsets such as T_H_2, T_H_17 and T_reg_ from the SILP as well as Tfh from Peyer’s patches showed reduced Ki67 expression (**Supplemental Figure 3B, C)**. Reduced Ki67 expression by ileal CD4 T cells was also observed in mice fed an SAA deficient diet compared to controls and was partially restored upon sulfide supplementation (**Fig 3G**). Further, the intensity of Nur77, a marker of T cell receptor engagement, was also significantly reduced in CD4 T cells following BSS treatment (**Fig 3H**). These results indicated an overall reduced fitness and activation of CD4 T cells following reduction in gut sulfides.

**Figure 3.**
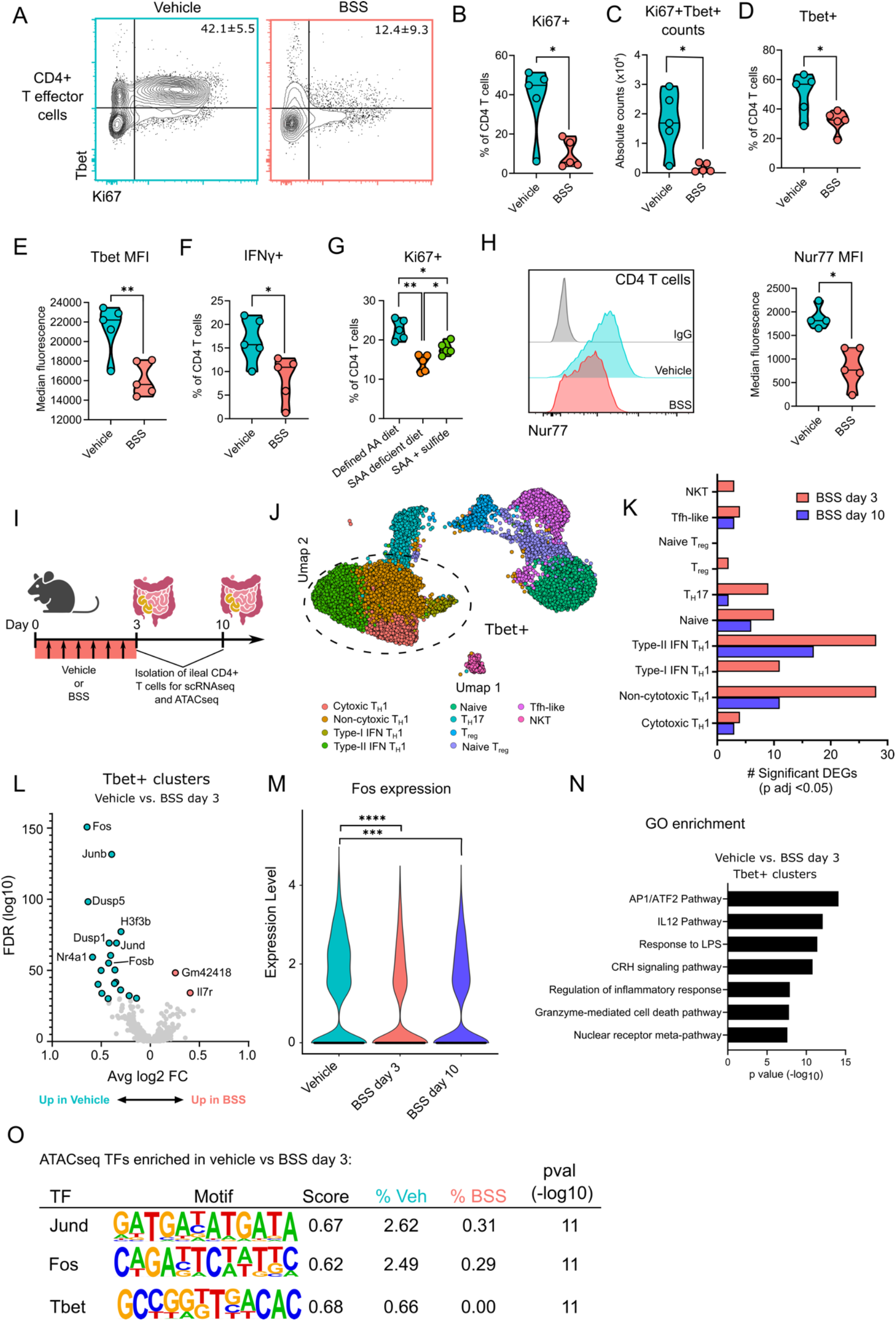
Gut CD4 T cell activation and AP-1 expression is impaired in the absence of gut sulfide. (**A**) Representative flow cytometry plots of CD4 effector T cells in SILP of vehicle and BSS treated mice at day 10 after start of treatment, with T-bet and Ki67 markers shown. **(B**) Frequency of Ki67+ CD4 T cells in SILP. (**C**) Total counts of Ki67+T-bet+ CD4 T cells in SILP. (**D**) Frequency of T-bet+ CD4 T cells in SILP. (**E**) Median fluorescence of T-bet in T-bet+ CD4 T cells in SILP. (**F**) Frequency of IFNγ+ CD4 T cells in SILP. (**G**) Frequency of Ki67+ CD4 T cells in SILP of mice on SAA diet or SAA diet + NaHS supplemented drinking water on day 21 of treatment. (**H**) Median fluorescence of Nur77 in SILP of vehicle or BSS treated mice CD4 T cells, with representative histograms shown, including isotype control (IgG). (**I**) Schematic of scRNAseq procedure, mice were treated with vehicle or BSS for 3 days and ileal CD4 T cells were isolated on day 3 or day 10 for scRNA sequencing and bulk ATAC seq. **(J)** UMAP of all CD4 T cells with identified clusters shown. Four T-bet+ T_H_1 clusters are noted with dotted oval. **(K**) Total number of significant differentially expressed genes in each cluster when comparing vehicle and BSS at day 3 or day 10, using Wilcox Rank Sum test. **(L**) Volcano plot of T-bet+ clusters comparing vehicle and BSS at day 3, with teal circles indicating significantly decreased and red significantly increased genes in BSS treated mice. (**M**) Violin plot of Fos expression among all CD4 T cells in vehicle, and days 3 and 10 of BSS treatment. **(N)** Gene ontogeny enrichment analysis of significantly upregulated genes in T-bet+ clusters, comparing vehicle vs BSS day 3. **(O)** Select ATAC-seq differentially enriched motifs, and their matching transcription factor identities, as analyzed by motif enrichment analysis of vehicle vs. BSS day 3 enriched peaks (TF, transcription factor). n.s. not significant, * p < 0.05, ** p < 0.01, *** p < 0.001, **** p < 0.0001, as measured by two-tailed t test (B-H) or Mann-Whitney U test (M).

To explore the underlying cause of impaired CD4 T cell activation, CD4 T cells were isolated from the ileum of vehicle or BSS treated mice before (day 3) and after (day 10) collapse to analyze their transcriptional profile as well as chromatin accessibility by scRNAseq and ATACseq, respectively (**Fig 3I**). Several clusters were identified, including four distinct T-bet^+^ T_H_1 clusters, two FoxP3^+^ T_reg_ clusters, Bcl6^+^ Tfh-like cells, RORγT^+^ T_H_17 cells, and CD62L^+^ naïve T cells (**Fig 3J**) (**Supplemental Figure 3D, 3E**). The four T_H_1 clusters were differentiated by their expression of cytotoxic markers such as granzyme and type I interferon stimulated genes (**Supplemental Figure 3E**). Aligned with our phenotypic observation (**Fig 1C**), differential gene expression analysis per cluster between vehicle treated and BSS treatment at day 3 or 10 revealed that the 4 T-bet expressing clusters were the most significantly impacted by BSS treatment, with the highest number of differentially expressed genes (DEGs) (**Fig 3K**). Further, there were more DEGs in each cluster at day 3 after BSS treatment compared to day 10. To understand how BSS treatment impacted T-bet expressing cells, we next combined T-bet^+^ clusters and quantified DEGs between vehicle and day 3 post BSS treatment. Several genes were downregulated in BSS treated T-bet+ CD4 T cells, including transcription factors *Fos, Junb* and *Jund*, dual specificity phosphatases *Dusp1* and *Dusp5*, and activation marker *Nr4a1* (Nur77) (**Fig 3L**). Of note, the transcription factor *Fos* was most significantly downregulated at both day 3 and day 10 post BSS treatment (**Fig 3M**). Fos and Jun proteins form the heterodimeric AP-1 transcription factor, which is a key modulator of CD4 T cell activation, proliferation and cytokine production (*30*). Gene ontology analysis of DEGs revealed significant enrichment in the AP-1 pathway and IL-12 pathway (**Fig 3N**). IL-12 pathway activation is known to be important for IFNγ production and T_H_1 function (*31*). We also quantified the expression of all predicted AP-1 targeted genes and observed that there was significant downregulation of this gene set in BSS treated T-bet^+^ cells (**Supplemental Figure 3F)**. This data supported the hypothesis that sulfide could augment activation of gut CD4 T cells by promoting expression of AP-1 transcription factors in T-bet expressing cells.

We next assessed the impact of BSS treatment on the epigenetic landscape of CD4 T cells, via ATACseq. To estimate changes in transcription factor binding, we performed motif enrichment analysis comparing regions of open chromatin unique to BSS treatment versus regions of open chromatin unique to vehicle. BSS treatment resulted in significantly reduced enrichment of AP-1 transcription factors motifs Jund and Fos, indicating reduced chromatin accessibility of their binding sites (**Fig 3O)**. Further, the T-bet motif was also reduced in the BSS treated condition compared to control (**Fig 3O**). These observations align with reduced T-bet expression in ileal CD4 T cells following BSS treatment (**Fig 3D, 3E)**. Taken together, bulk ATAC-seq and scRNAseq analysis supported the hypothesis that reduced AP-1 activity following sulfide depletion could contribute to reduced CD4 T cell fitness within the gut.

### Sulfide acts directly on CD4 T cells to promote proliferation and activation

We next explored the potential mechanism by which sulfide could be modulating AP-1 activity in CD4 T cells. To this end, primary naïve CD4 T cells were isolated and then stimulated *in vitro* prior to exposure to media containing a slow releasing sulfide donor GYY4137 (**Fig 4A**). In agreement with our *in vivo* data (**Fig 1C**), CD4 T cells grown in the presence of sulfide *in vitro* expanded to greater numbers than those without sulfide after 5 days of culture (**Fig 4B**). Increased cell number was associated with an increased rate of proliferation, as evidenced by elevated Ki67 expression and increased cell division as measured by dilution of cell tracing dye (**Fig 4C, 4D**). Importantly, there was no significant decrease in cell death in sulfide-cultured cells, supporting the idea that increased cell number was due to increased proliferation rather than reduced cell death (**Supplemental Figure 4**). Sulfide-exposed CD4 T cells also showed enhanced activation *in vitro*, as measured by Nur77 MFI (**Fig 4E**). Thus, sulfide exposure *in vitro* had positive impact on T cell proliferation and activation, further supporting our hypothesis that gut sulfides promote CD4 T cell fitness within the gut.

**Figure 4.**
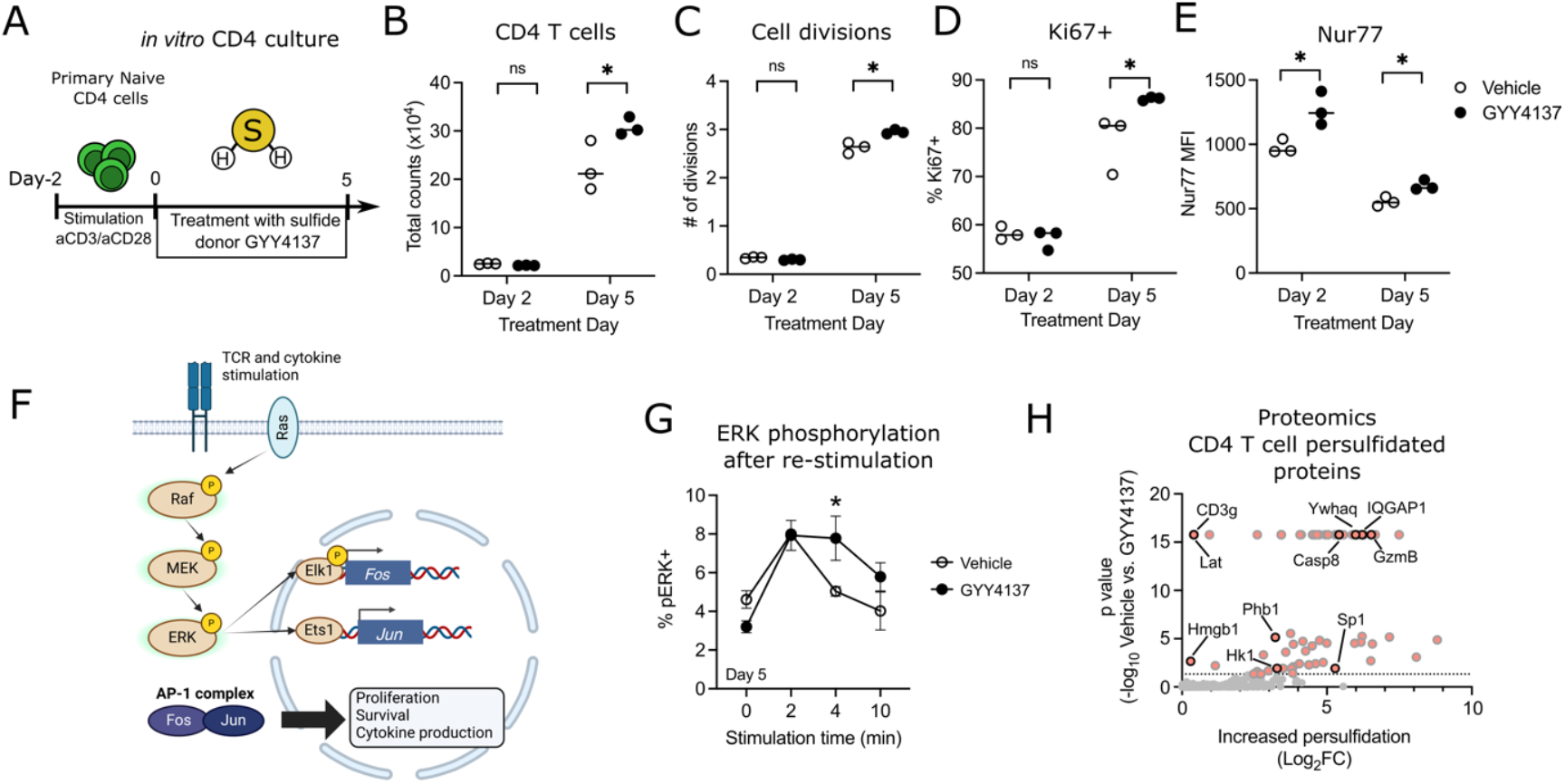
Sulfide acts directly on CD4 T cells to modulate MAP kinase phosphorylation. **(A)** *In vitro* culture procedure, naïve CD4 T cells isolated from splenocytes and lymph nodes were cultured for 2 days with plate bound αCD3/αCD28. Cells were then moved to new wells with or without 100μM sulfide donor GYY4137 for 5 days without stimulation. **(B)** Total counts of CD4 T cells at days 2 and 5 of sulfide treatment. **(C)** Mean number of cell divisions for CD4 T cells at days 2 and 5 of sulfide treatment. **(D**,**E)** CD4 T cells at day 2 and 5 of sulfide treatment assessed for percent Ki67+ (D) or Nur77 median fluorescence (E). **(F)** TCR signaling leading to *Fos* and *Jun* transcription via MAPK cascade. **(G)** CD4 T cells cultured for 5 days with sulfide were restimulated with crosslinked αCD3 for 0-10 minutes and assessed for Erk phosphorylation by flow cytometry. Phospho-ERK+ cells shown as defined by isotype control antibody. **(H)** CD4 T cells treated with GYY4137 for 3 days were subjected to the ProPerDP protocol to enrich for persulfidated proteins. Volcano plot shows significantly enriched proteins in GYY4137 treated cells, with proteins of interest highlighted. * p < 0.05, as measured by two-tailed t test

Transcriptional analysis pointed to a reduction in AP-1 transcription in the absence of sulfide (**Fig 3N**). Previous work established that transcription of AP-1 subunits *Fos* and *Jun* is driven by the multifunctional MAP kinase pathway which terminates in Erk phosphorylation activating several downstream transcription factors (**Fig 4F**) (*32*). To assess whether Erk phosphorylation was modified in sulfide treated CD4 T cells, we assessed the level of Erk phosphorylation of restimulated CD4 T cells cultured in sulfide as in Fig 4A. While sulfide did not alter peak Erk phosphorylation, Erk maintained phosphorylation for longer in sulfide exposed T cells than in untreated cells (**Fig 4F**). This modest increase in Erk phosphorylation duration could potentially contribute to the observed increased AP-1 transcription in sulfide exposed gut CD4 T cells.

While sulfide is a promiscuous molecule, most of its activity is thought to be mediated by its ability to provide a specific post-translational modification to proteins (*33*). Free thiol groups from cysteine residues in proteins can be converted to a reactive persulfide residue, which can alter the function of select proteins (*34*). To interrogate whether sulfide post-translationally modified CD4 T cell proteins, we used a proteomics-based method to enrich for persulfide containing proteins (*35*). Proteomics revealed 91 proteins with significantly increased persulfidation following *in vitro* sulfide treatment (**Fig 4G, Table S2)**. This included TCR signaling proteins Lat and CD3γ, cell death protein caspase 8, cytotoxic protein granzyme B, and phospho binding protein 14-3-3 theta (Ywhaq). A top hit was scaffolding protein IQGAP1, previously shown to enhance the activity of the MAPK cascade (*36*). These data indicated that sulfide could act directly on CD4 T cells to promote their activation and resulted in persulfide modification of specific proteins.

### Gut sulfides are required for T-dependent antibody response to oral vaccination

Because of the significant impact of BSS on Peyer’s patch T and B cells, we assessed the potential role of sulfide in oral immunization response. To this end, we depleted gut sulfides with BSS and subsequently orally vaccinated mice with cholera toxin (CTX) and ovalbumin (OVA). In this model, CTX acts as an adjuvant while both CTX and OVA are the novel antigens (**Fig 5A**). In mice orally vaccinated with CTX+OVA and treated with BSS, the number of CD4 T cells in Peyer’s Patches was significantly lower compared to vaccinated mice treated with the vehicle (**Fig 4B**). This reduction was accompanied by decreased proliferation, as indicated by lower Ki67 expression (**Fig 5C)**. Tfh cells and GC B cells were also significantly decreased in BSS treated mice after vaccination, indicative of a reduced germinal center response (**Fig 5D, 5E**). Vaccination in vehicle treated mice induced a robust immune response associated with CTX specific IgA secreting cells found in the ileum (**Fig 5F, 5G**). On the other hand, mice vaccinated and treated with BSS displayed a significant reduction in antibody response with fewer antigen-specific IgA secreting cells (**Fig 5F, 5G**). To track the antigen specific T cell response, we transferred OVA-specific OT-II cells prior to vaccination. The absolute number of OT-II cells in the ileum was significantly decreased after sulfide depletion in vaccinated mice (**Fig 5H**).

**Figure 5.**
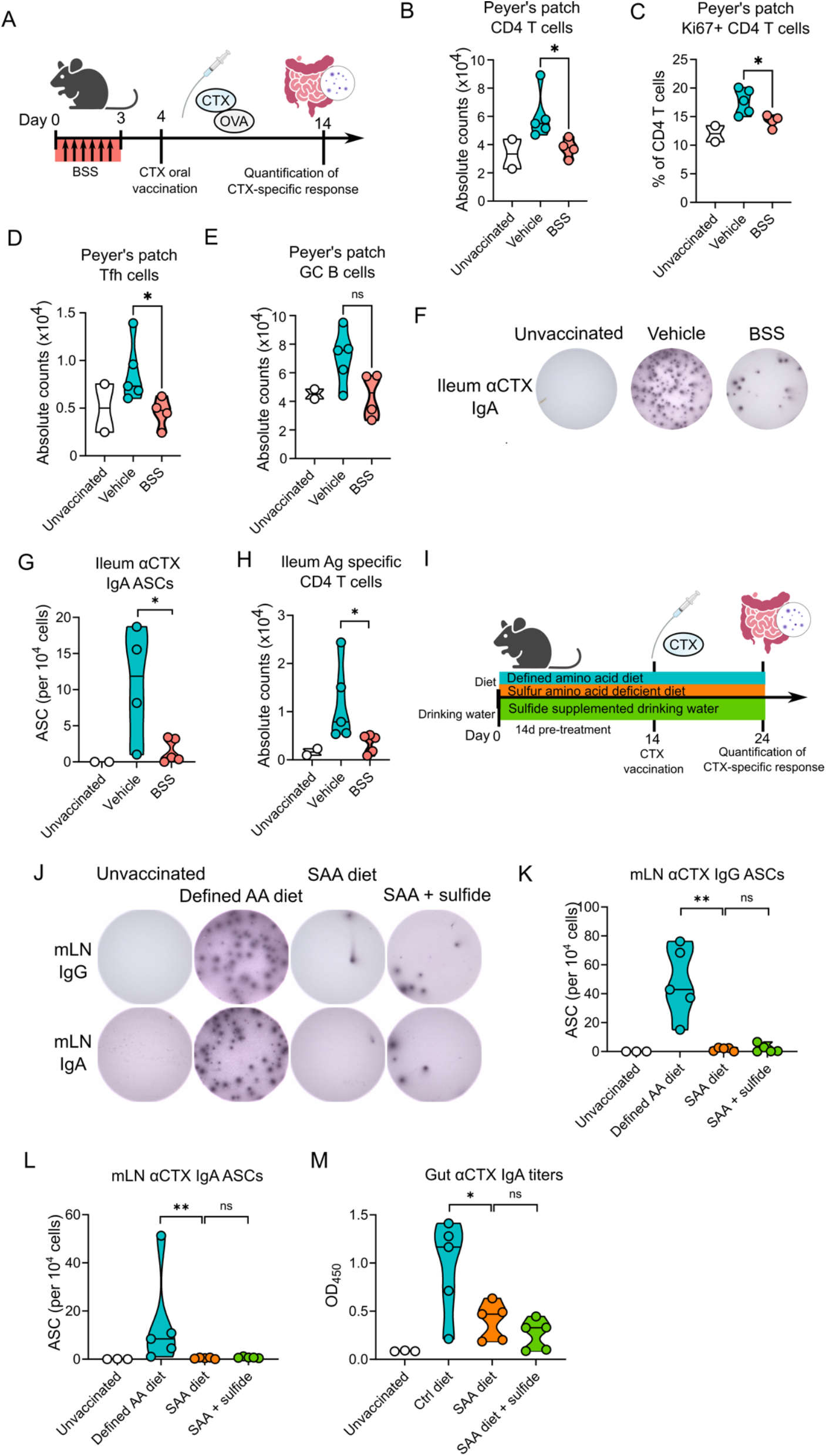
Gut sulfides are required for productive mucosal vaccine response. **(A)** Vaccination of BSS treated mice with oral gavage of cholera toxin and ovalbumin one day after 3 days of BSS treatment, with vaccine response analyzed 10 days later. (**B**) Absolute counts of Peyer’s patch CD4 T cells in unvaccinated mice, or vaccinated mice treated with vehicle or BSS. (**C**) Frequency of Ki67+ CD4 T cells in Peyer’s patches. (**D**,**E**) Absolute counts of Tfh cells (D) and GC B cells (E) in Peyer’s patches. **(F, G)** ELISPOT of cholera toxin specific IgA secreting cells in the ileum of vaccinated mice, as shown by representative ELISPOT wells (F) and total frequency of antibody secreting cells (G). **(H)** Total counts of OVA specific transferred OT-II cells in ileum of vaccinated mice. **(I)** Vaccination of SAA diet fed mice after 2 weeks on SAA or defined AA diet, then assessed for vaccine response 10 days later. **(J-L)** ELISPOT of cholera toxin specific IgG and IgA secreting cells in the mLN of vaccinated mice, as shown by representative ELISPOT wells (J) and total frequency in mLN for cholera toxin specific IgG (K) and IgA secreting cells (L). **(M)** Cholera toxin specific gut IgA titers as measured by ELISA of gut luminal content. n.s. not significant, * p < 0.05, ** p < 0.01, *** p < 0.001, as measured by two-tailed t test.

To mimic a setting of reduced sulfur amino acid intake in the context of mucosal vaccination we explored mucosal vaccine response using SAA dietary restriction, vaccinating mice after 2 weeks on an SAA restricted diet (**Fig 5I**). Overall vaccine response was abrogated in SAA restricted diet, a reduction significantly stronger than that following BSS sulfide depletion. We observed a significant defect in the expansion of germinal center cells in the Peyer’s patch after vaccination in mice on SAA diet (**Supplemental Figure 5A-C**). Strikingly, there was a near complete loss of specific antibody secreting cells in the draining lymph node (**Fig 5J**). While there were robust IgA and IgG secreting cells in response to CTX on the control diet, there was no significant response in mice on an SAA diet, for both IgA (**Fig 5K**) and IgG (**Fig 5L**) in the draining lymph node. This translated into significantly reduced antibody production in the gut, with reduced αCTX IgA found secreted into the gut lumen (**Fig 5M**). This antibody response is the major correlate of protection in many mucosal vaccines (*37*). These data highlight the central and previously unappreciated role of gut sulfides in mediating mucosal vaccine response, providing a mechanistic hypothesis for the known reduced response to oral vaccine in the context of malnutrition and more specifically low protein diet (*38*).

## Discussion

The role of endogenous hydrogen sulfide in eukaryotes has only recently been appreciated, but its relevance in numerous biological processes is now well established. Here we show a novel role for hydrogen sulfide in a region where it is most densely concentrated in the gastrointestinal tract. Key cell types that mediate mucosal immunity were affected including T helper 1 cells and T follicular helper cells. T_H_1 cells are key producers of interferon gamma and have been implicated in protection against a variety of infections. This includes several enteric bacterial infections, such as *Salmonella* and *Helicobacter pylori* (*39, 40*). A loss of T_H_1 cells in the absence of sulfide suggests that gut sulfides may play a critical role in protection against infection (*28*). The loss of T follicular helper cells reflects a key defect in the generation of a germinal center response, which is reflected in the severe impairment of mucosal vaccine responses in the absence of sulfide.

Several other studies have given evidence that CD4 T cells are impacted by endogenous sulfide. Ou et al show that T_H_17 cell differentiation is inhibited by apoptosis induced sulfide production in spleen and blood (*41*). CD4^+^ T regulatory cell differentiation has been reported to be enhanced by serum sulfide concentrations, providing a role for sulfide to protect against autoimmunity (*42*). Ji et al observed that in a methionine-restricted diet, mice were more susceptible to gastrointestinal cancers, due to a reduction in systemic T cells (*43*). They additionally observed lower IFNγ RNA in intestinal tissue, echoing our findings in SILP T_H_1 cells (**Fig 3F**).

In this study, we propose that sulfide acts directly on CD4 T cells, enhancing their activation and proliferation within the gut. In the absence of this sulfide signal, we observed an acute contraction of CD4 T cells. As it is difficult to assess cell death *in vivo*, it is unclear whether this loss is being driven by increased cell death in the absence of sulfide. However, *in vitro* exposure to sulfide did not alter CD4 T cell death (**Supplemental Figure 4A**). The enhanced activation of sulfide exposed CD4 T cells was associated with an augmentation of AP-1 expression, which is regulated by activation of the MAPK cascade. Our data also suggests that sulfide could augment phosphorylation of MAPK protein Erk, though additional mechanistic experiments would be required to confirm this pathway. Erk phosphorylation has previously been shown to be enhanced after sulfide exposure in an alternate cell type, kidney tubule epithelial cells (*44*). Phospho-Erk was also shown to be upregulated in the context of sepsis by endogenous sulfide production in the lungs and liver (*45*). Thus, sulfide augmented Erk signaling may be a generalizable phenomenon in many cell types. Erk is a key signaling molecule in several T cell activation pathways, including TCR engagement, costimulation and cytokine signaling (*46, 47*). Any alterations to the phosphorylation status of Erk would significantly alter CD4 T cell function. We also observed alterations to chromatin accessibility of T-bet, the main transcription factor in T_H_1 cells which are most impacted by sulfide in the gut. It is unclear why T_H_1 cells are most affected, or if sulfide impacts CD4 T cell polarization. Each CD4 T cell subset warrants further investigation into possible distinct interactions with gut sulfides.

Sulfide has no known conventional receptor, importer or sensor within cells, instead it readily diffuses through the membrane and alters protein structure (*33*). Free cysteine thiols are modified to persulfide groups, which creates a more reactive cysteine residue and can alter protein function (*34*). Proteomics of the persulfide enriched protein fraction in CD4 T cells revealed many possible targets of sulfide which could lead to observed changes in Erk phosphorylation. A top hit IQGAP1 is a scaffolding protein, which previously has been shown to enhance the function of the Erk-MAPK pathway (*36*). Ninety-one proteins were enriched for persulfide residues, so it is possible that several proteins may have altered function that led to the observed CD4 phenotype. It will be critical to explore how this “persulfidome” regulates CD4 T cell biology.

Our bismuth-based model for depletion of gut sulfides specifically targeted only gut sulfides, as it is not absorbed systemically and should not directly alter systemic sulfide levels. We designed the protocol to approximate human usage of this common over-the-counter drug. As we observed significant impacts through normal use of this drug in mice, it warrants further investigation into how this could translate into humans. Drugs that disrupt gut sulfides could adversely affect mucosal immune function, or impact susceptibility to enteric infections. Our diet-based model of sulfide modulation also has translational implications in humans. It is known that dietary intake of protein can affect the amount of sulfide produced in the human gut (*14*). People who primarily eat plant-based protein, or who have low protein diets, may have lower gut sulfide concentrations (*48*). How this could impact mucosal immunity in these populations has yet to be explored. In the context of mucosal vaccination, the influence diet driven variation in gut sulfides is a key question. Mucosal vaccination is often employed in low-resource areas, where hospital facilities are sparse and field-based vaccination is more prudent (*49*). Unfortunately, oral vaccines have proven challenging to develop, with noted lack of immunogenicity and variations in response across populations (*50*). In many of these resource poor areas, malnutrition and in particular low protein intake, is a common concern. These low protein diets may lead to reduced gut sulfides at the time of mucosal vaccination. Our study suggests that this could lower efficacy of vaccination in these populations. Complementation of gut sulfides, possibly through a sulfide donating molecule in these vaccines, may lead to increased efficacy of oral vaccination in these populations. Gut targeted sulfide donors have been developed which could be candidates for this strategy (*51*). Overall, it is clear from this study that gut sulfides play a central role in gut immune homeostasis **(Supplemental Figure 5D**), warranting their consideration as a marker of gut health.

## Notes

## Acknowledgements

This work was supported by the National Institute of Allergy and Infectious Diseases (NIAID) of the National Institutes of Health under the Division of Intramural Research, NIAID, NIH. The authors would like to thank V. Link and A. Wells for critical reading of the manuscript and editorial input; the NIAID flow cytometry unit for cell sorting assistance; the NIAID Gnotobiotic Mouse Facility for germ free experiments; E. Lewis and K. Beacht for technical support. This work utilized the computational resources of the NIAID HPC Locus and Skyline clusters. Schematic figures were created with BioRender (https://biorender.com).

## Funding

National Institutes of Health Division of Intramural Research of NIAID grant 1ZIA-AI001115 and 1ZIA-AI001132 (VIB, IG, AS, NB, JC, ME, MN, YB)

NIAID Division of Intramural Research (SH, DG, PLS, VB, ANL, SM, AB, PJPC) National Institutes of Health Office of Dietary Supplements (VIB)

CRI Irvington Postdoctoral Fellowship (ME) NIGMS Prat Fellowship (AS)

## Author contributions

Conceptualization: VIB, YB

Methodology: VIB, VB, IG, SM, AB, PJPC, ME, DG

Validation: VIB

Formal analysis: VIB, IG, JC, VB

Investigation: VIB, IG, AS, NB, MN, ME, VB, DG, SH

Visualization: VIB, IG

Funding acquisition: YB, ANL, SH, PLS Project administration: YB

Supervision: YB, ANL, SH, PLS Writing – original draft: VIB, VB

Writing – review & editing: VIB, YB, SH

## Data and materials availability

All data are available in the main text or supplemental figures. Raw genomic data from single-cell RNAseq, ATAC-seq, 16S rDNA sequencing, and metagenomics is deposited on NCBI and proteomics data is deposited on PRIDE (PXD061194).

## Supplementary Materials

Materials and Methods

Supplementary Figs. S1 to S5

Tables S1 to S3

## References

1. E. Cuevasanta, M. N. Moller, B. Alvarez, Biological chemistry of hydrogen sulfide and persulfides. Arch Biochem Biophys 617, 9–25 (2017).

2. K. R. Olson, K. D. Straub, The Role of Hydrogen Sulfide in Evolution and the Evolution of Hydrogen Sulfide in Metabolism and Signaling. Physiology (Bethesda) 31, 60–72 (2016).

3. S. Ranjan, Z. R. Todd, J. D. Sutherland, D. D. Sasselov, Sulfidic Anion Concentrations on Early Earth for Surficial Origins-of-Life Chemistry. Astrobiology 18, 1023–1040 (2018).

4. S. Han, Y. Li, H. Gao, Generation and Physiology of Hydrogen Sulfide and Reactive Sulfur Species in Bacteria. Antioxidants (Basel) 11, (2022).

5. R. Konrad et al., Distribution and Activity of Sulfur-Metabolizing Bacteria along the Temperature Gradient in Phototrophic Mats of the Chilean Hot Spring Porcelana. Microorganisms 11, (2023).

6. G. Cirino, C. Szabo, A. Papapetropoulos, Physiological roles of hydrogen sulfide in mammalian cells, tissues, and organs. Physiol Rev 103, 31–276 (2023).

7. X. L. He et al., Hydrogen sulfide improves spatial memory impairment and decreases production of Abeta in APP/PS1 transgenic mice. Neurochem Int 67, 1–8 (2014).

8. W. R. Dunn, S. P. Alexander, V. Ralevic, R. E. Roberts, Effects of hydrogen sulphide in smooth muscle. Pharmacol Ther 158, 101–113 (2016).

9. M. Whiteman, P. G. Winyard, Hydrogen sulfide and inflammation: the good, the bad, the ugly and the promising. Expert Rev Clin Pharmacol 4, 13–32 (2011).

10. J. Furne, A. Saeed, M. D. Levitt, Whole tissue hydrogen sulfide concentrations are orders of magnitude lower than presently accepted values. Am J Physiol Regul Integr Comp Physiol 295, R1479–1485 (2008).

11. J. Jiang et al., Hydrogen Sulfide--Mechanisms of Toxicity and Development of an Antidote. Sci Rep 6, 20831 (2016).

12. D. Dordevic, S. Jancikova, M. Vitezova, I. Kushkevych, Hydrogen sulfide toxicity in the gut environment: Meta-analysis of sulfate-reducing and lactic acid bacteria in inflammatory processes. J Adv Res 27, 55–69 (2021).

13. K. L. Flannigan, K. D. McCoy, J. L. Wallace, Eukaryotic and prokaryotic contributions to colonic hydrogen sulfide synthesis. Am J Physiol Gastrointest Liver Physiol 301, G188–193 (2011).

14. E. A. Magee, C. J. Richardson, R. Hughes, J. H. Cummings, Contribution of dietary protein to sulfide production in the large intestine: an in vitro and a controlled feeding study in humans. Am J Clin Nutr 72, 1488–1494 (2000).

15. M. C. Fitzpatrick, A. V. Kurpad, C. P. Duggan, S. Ghosh, D. G. Maxwell, Dietary intake of sulfur amino acids and risk of kwashiorkor malnutrition in eastern Democratic Republic of the Congo. Am J Clin Nutr 114, 925–933 (2021).

16. F. L. Suarez, J. K. Furne, J. Springfield, M. D. Levitt, Bismuth subsalicylate markedly decreases hydrogen sulfide release in the human colon. Gastroenterology 114, 923–929 (1998).

17. M. C. Pitcher, E. R. Beatty, J. H. Cummings, The contribution of sulphate reducing bacteria and 5-aminosalicylic acid to faecal sulphide in patients with ulcerative colitis. Gut 46, 64–72 (2000).

18. G. A. Castro, C. J. Arntzen, Immunophysiology of the gut: a research frontier for integrative studies of the common mucosal immune system. Am J Physiol 265, G599–610 (1993).

19. B. A. H. Jensen et al., Small intestine vs. colon ecology and physiology: Why it matters in probiotic administration. Cell Rep Med 4, 101190 (2023).

20. A. B. Nair, S. Jacob, A simple practice guide for dose conversion between animals and human. J Basic Clin Pharm 7, 27–31 (2016).

21. H. J. Van Kruiningen, A. B. West, B. J. Freda, K. A. Holmes, Distribution of Peyer’s patches in the distal ileum. Inflamm Bowel Dis 8, 180–185 (2002).

22. M. A. Mintz, J. G. Cyster, T follicular helper cells in germinal center B cell selection and lymphomagenesis. Immunol Rev 296, 48–61 (2020).

23. A. G. Buret, T. Allain, J. P. Motta, J. L. Wallace, Effects of Hydrogen Sulfide on the Microbiome: From Toxicity to Therapy. Antioxid Redox Signal 36, 211–219 (2022).

24. A. Ghasemi, S. Jeddi, N. Yousefzadeh, K. Kashfi, R. Norouzirad, Dissolving sodium hydrosulfide in drinking water is not a good source of hydrogen sulfide for animal studies. Sci Rep 13, 21839 (2023).

25. Y. Belkaid, T. W. Hand, Role of the microbiota in immunity and inflammation. Cell 157, 121–141 (2014).

26. J. M. Wells, Immunomodulatory mechanisms of lactobacilli. Microb Cell Fact 10 Suppl 1, S17 (2011).

27. Ivanov, II et al., Induction of intestinal Th17 cells by segmented filamentous bacteria. Cell 139, 485–498 (2009).

28. A. Stacy et al., Infection trains the host for microbiota-enhanced resistance to pathogens. Cell 184, 615–627 e617 (2021).

29. F. Faber et al., Host-mediated sugar oxidation promotes post-antibiotic pathogen expansion. Nature 534, 697–699 (2016).

30. M. Yukawa et al., AP-1 activity induced by co-stimulation is required for chromatin opening during T cell activation. J Exp Med 217, (2020).

31. K. A. Ullrich et al., Immunology of IL-12: An update on functional activities and implications for disease. EXCLI J 19, 1563–1589 (2020).

32. M. Rincon, R. A. Flavell, R. J. Davis, Signal transduction by MAP kinases in T lymphocytes. Oncogene 20, 2490–2497 (2001).

33. T. Vignane, M. R. Filipovic, Emerging Chemical Biology of Protein Persulfidation. Antioxid Redox Signal 39, 19–39 (2023).

34. C. T. Yang, N. O. Devarie-Baez, A. Hamsath, X. D. Fu, M. Xian, S-Persulfidation: Chemistry, Chemical Biology, and Significance in Health and Disease. Antioxid Redox Signal 33, 1092–1114 (2020).

35. E. Doka et al., A novel persulfide detection method reveals protein persulfide-and polysulfide-reducing functions of thioredoxin and glutathione systems. Sci Adv 2, e1500968 (2016).

36. M. Roy, Z. Li, D. B. Sacks, IQGAP1 is a scaffold for mitogen-activated protein kinase signaling. Mol Cell Biol 25, 7940–7952 (2005).

37. M. Gagne et al., Mucosal adenovirus vaccine boosting elicits IgA and durably prevents XBB.1.16 infection in nonhuman primates. Nat Immunol 25, 1913–1927 (2024).

38. S. Rho et al., Protein energy malnutrition alters mucosal IgA responses and reduces mucosal vaccine efficacy in mice. Immunol Lett 190, 247–256 (2017).

39. N. Bagheri, L. Salimzadeh, H. Shirzad, The role of T helper 1-cell response in Helicobacter pylori-infection. Microb Pathog 123, 1–8 (2018).

40. R. Ravindran, S. J. McSorley, Tracking the dynamics of T-cell activation in response to Salmonella infection. Immunology 114, 450–458 (2005).

41. Q. Ou et al., Apoptosis releases hydrogen sulfide to inhibit Th17 cell differentiation. Cell Metab 36, 78–89 e75 (2024).

42. R. Yang et al., Hydrogen Sulfide Promotes Tet1- and Tet2-Mediated Foxp3 Demethylation to Drive Regulatory T Cell Differentiation and Maintain Immune Homeostasis. Immunity 43, 251–263 (2015).

43. M. Ji et al., Methionine restriction-induced sulfur deficiency impairs antitumour immunity partially through gut microbiota. Nat Metab 5, 1526–1543 (2023).

44. S. J. Han, J. I. Kim, J. H. Lipschutz, K. M. Park, Hydrogen sulfide, a gaseous signaling molecule, elongates primary cilia on kidney tubular epithelial cells by activating extracellular signal-regulated kinase. Korean J Physiol Pharmacol 25, 593–601 (2021).

45. H. Zhang, S. M. Moochhala, M. Bhatia, Endogenous hydrogen sulfide regulates inflammatory response by activating the ERK pathway in polymicrobial sepsis. J Immunol 181, 4320–4331 (2008).

46. C. F. Chang et al., Polar opposites: Erk direction of CD4 T cell subsets. J Immunol 189, 721–731 (2012).

47. K. Adachi, M. M. Davis, T-cell receptor ligation induces distinct signaling pathways in naive vs. antigen-experienced T cells. Proc Natl Acad Sci U S A 108, 1549–1554 (2011).

48. R. Siener, N. Bitterlich, H. Birwe, A. Hesse, The Impact of Diet on Urinary Risk Factors for Cystine Stone Formation. Nutrients 13, (2021).

49. T. Azegami, Y. Yuki, H. Kiyono, Challenges in mucosal vaccines for the control of infectious diseases. Int Immunol 26, 517–528 (2014).

50. M. M. Levine, Immunogenicity and efficacy of oral vaccines in developing countries: lessons from a live cholera vaccine. BMC Biol 8, 129 (2010).

51. K. M. Dillon et al., Targeted Delivery of Persulfides to the Gut: Effects on the Microbiome. Angew Chem Int Ed Engl 60, 6061–6067 (2021).

