## Supplemental Text for "Sulfide is a keystone metabolite for gut homeostasis and immunity"

### Materials and Methods

Mice

All wild-type mice used were C57BL6/J female and male mice aged 8-14 weeks at the start of the experiment, and were sex and age matched. For cholera vaccination experiments, T cells were isolated from Tg(Tcratcrb)425cbn,Rag1<tm1Mom> (OT-II Transgenic CD45.2+) mice and transferred into C57/Bl6J x B6.SJL-CD45a(Ly5a)Nai F1 (CD45.1/CD45.2+). All mice were obtained from the Taconic-NIAID exchange program. For all experiments, littermate control mice were randomized into cages for 1 week before beginning experimental manipulations to correct for cage effect. Specific pathogen free (SPF) mice were bred and maintained under SPF conditions at an American Association for the Accreditation of Laboratory Animal Care (AAALAC)–accredited animal facility at NIAID and housed in accordance with the procedures outlined in the Guide for the Care and Use of Laboratory Animals. Germ-free and gnotobiotic colonized mice were housed in the NIAID Gnotobiotic Animal Facility under gnotobiotic conditions, and were all bred in the facility. All experiments performed under an approved animal study proposal (LHIM2E), approved by the NIAID IACUC.

Bismuth subsalicylate treatment

Mice were treated with BSS by oral gavage or via drinking water. Mice treated by gavage were given 5mg BSS in H2O orally by gavage, every 12 hours for 72 hours, for a total of 6 doses. Mice treated in drinking water were put on special drinking water containing 7mg/mL BSS for 72 hours. Precipitated BSS was resolubilized by shaking every 24 hours. Germ-free BSS treatment was done in drinking water as noted, and BSS containing drinking water was autoclaved before use. Vehicle treated mice were given $H_2O$ at identical doses and times.

SAA deficient diet

Mice were placed on an SAA deficient diet containing no protein, and defined amounts of amino acids. The SAA diet contained 0.1% methionine and no cysteine/cystine, 66% carbohydrate, while the control AA diet contained 0.82% methionine and 0.35% cystine, 64.9% carbohydrates. All diets contain 8% fat and were irradiated. Mice were switched from standard chow to AA diets for 2 weeks. Some mice on SAA diet were also supplemented with sulfide drinking water, containing 0.1 mg/mL sodium hydrosulfide, with pH adjusted to 8.0, which was replaced daily.

Murine tissue processing

Intestinal tissue lymphocytes were isolated as previously described (*1*). Briefly, small and large intestine were removed from mice, with large intestinal tissue including the cecum. For subsections of the small intestine, the tissue length was measured and split into even thirds for duodenum, jejunum and ileum samples. Epithelium was dissociated in T cell media (RPMI 1640 media supplemented with 2 mM L-glutamine, 1 mM sodium pyruvate and nonessential amino acids, 55 mM β-mercaptoethanol, 20 mM HEPES, 100 U/ml penicillin, 100 μg/ml streptomycin) plus 3% FBS, 100ug/mL dithiothreitol and 5mM EDTA for 20 minutes at 37°C 5% $CO_2$, and then washed 3 times with dissociation media (T cell media with 5mM EDTA). The dissociated fraction was used for intraepithelial lymphocytes. The remaining tissue was minced and digested for 30 min in digest media (T cell media with 100 μg/ml liberase TI

(Roche) and 500 μg/ml DNase I (Sigma-Aldrich)) and filtered with a 70μm filter. Both fractions were then separated with 37.5% Percoll to isolate lymphocytes. Lymph nodes and Peyer's patches were smashed through a 70μm filter and centrifuged to isolate lymphocytes.

Sulfide measurements

Free sulfides in the gut were measured using an amperometric microsensor (Type-I, Unisense). Values were calibrated with sodium hydrosulfide stock solutions. Free sulfide was measured in situ by piercing the gut and inserting the sensor fully into the lumen, recording measurements after 10 seconds of stabilization. Fecal sulfide levels were measured using the Oxiselect free hydrogen sulfide kit (Cell Biolabs). Fecal pellets were homogenized in H2O with 0.1N NaOH to prevent release of sulfide during sample preparation. Supernatant was added to 0.2N HCl to lower pH and release sulfide during measurements.

Microbiome sequencing

Samples were collected fresh from mice and kept at -20ºC until sequencing. Fecal samples were collected from live mice in sterile collection cups. SI luminal content was collected at necropsy by removing tissue and shaking vigorously in PBS to dissociate luminal bacteria. Mucosal associated bacteria were then isolated by scraping this tissue with forceps to remove mucus layer and shaken vigorously in PBS. Fractions were then spun down at 5000xg for 10 min to pellet bacteria. Fecal DNA was purified using the MagAttract PowerMicrobiome DNA/RNA kit (Qiagen). Amplification of the V4 hypervariable region of the bacterial 16*S* rRNA gene was performed using the 515f and 806r primers (515F: 5′ -GTGCCAGCMGCCGCGGTAA-3′; 806R: 5′ -GGACTACHVGGGTWTCTAAT-3′), followed by an additional PCR to append unique barcodes to each sample. Amplicons were quantified using Kapa Library Quantification Complete Kit (ROX Low) (Kapa Biosystems) and pooled at equimolar concentrations before being sequenced on Illumina NexSeq or MiSeq. Metagenomic libraries were constructed from 100 ng of DNA as starting material using the Illumina DNA Prep kit. Illumina DNA/RNA UD Indexes were used to add sample-specific sequencing indices to both ends of the libraries. An Agilent 4200 TapeStation system with High Sensitivity D5000 ScreenTape (Agilent Technologies, Inc) was used to verify quality and assess final library size. Metagenomic and 16S libraries were normalized and pooled at an equimolar concentration. Final pools were diluted to 750 pM and sequenced on a NextSeq2000 sequencer using a paired-end (100x100) NextSeq 1000/2000 P2 (200 cycles) kit (Illumina, Inc). 16S sequencing data was analyzed with QIIME2 (*2*) using the dada2 denoising algorithm to generate ASVs and the Greengenes 13.8 database for taxonomic classification. Metagenomic data was analyzed with the Humann 3 pipeline (*3*).

Single cell RNA sequencing and ATAC-Seq

Single cell RNA sequencing was done on sorted CD4 T cells. Single cell suspensions of ileal lymphocytes were labelled with Total-Seq C hashtag oligonucleotide antibodies and antibodies against surface markers, then sorted on a Sony SH800S sorter for live CD45+CD90+TCRB+ CD4+CD8- cells. Samples were pooled and loaded onto a 10X Chromium Single Cell Controller, and libraries prepared using the 10X Next GEM 5' kit (10X Genomics). Libraries were sequenced with on an Illumina NexSeq 1000 sequencer. Sequencing data was analyzed with Cellranger (10X genomics) and Seurat (*4*). ATAC-Seq was performed as previously described (*5*). Briefly, cells were lysed with lysis buffer (0.1% NP-40, 0.1% Tween 20, 0.01% Digitonin) washed, and centrifuged to isolate nuclei. Transposition was performed with Illumina Tagment DNA enzyme TDE1 and total DNA was then purified with a MinElute Reaction Cleanup Kit (QIAGEN). Library was PCR amplified using Illumina/Nextera i5 primers (Ad1 5'- AATGATACGGCGACCACCGAGATCTACACTCGTCGGCAGCGTCAGATGTG-3', Ad2 5'- CAAGCAGAAGACGGCATACGAGATxxxxxxxx GTCTCGTGGGCTCGGAGATGT-3', where x represents i7 index adapter). Libraries were sequenced on an Illumina NexSeq 1000 sequencer. Reads were trimmed with trimmomatic (*6*), aligned with STAR (*7*), and peaks annotated using Homer (*8*). Differential peak analysis was performed with the Homer getDiffPeaksReplicates command, with BSS treated group used as background peaks and significant motifs reported.

<u>Flow cytometry</u>

Single cell suspensions were stained with cocktails of fluorophore-conjugated antibodies, as listed in Table S3. For intracellular staining, cells were fixed with Foxp3/Transcription Factor Staining Buffer Set (eBioscience) and stained with fluorophore-conjugated antibodies for at least 60 min at 4°C. Intracellular cytokine staining was performed on cells that were stimulated for 150 minutes in media containing phorbol myristate (50ng/mL), 5mg/mL ionomycin and a 1:1000 dilution of GolgiPlug (BD Biosciences). All staining was performed using purified anti-mouse CD16/32 and purified rat gamma globulin (Jackson ImmunoResearch) as a blocker of Fc interactions. Cells were analyzed on an LSR Fortessa (BD) running FACS Diva software. Data was analyzed using FlowJo software.

<u>Confocal microscopy</u>

Small intestine, Peyer's patches and lymph nodes were fixed with 10 mg/ml paraformaldehyde for 24 h. Samples were washed in phosphate buffer and dehydrated in 30% sucrose phosphate buffer, mounted in O.C.T. Compound (Fisher) and snap-frozen on dry ice. Sections with 11 µm thickness were cut on a cryostat. Sections were washed in phosphate buffer and blocked in 1% BSA, 0.25% Triton X blocking buffer for 30 min at room temperature. Tissues were stained overnight at 4 °C in blocking buffer with antibodies (**Table S3**) overnight at 4°C. After being washed three times with PBS, tissues were mounted with ProLong Gold (Molecular Probes) antifade reagent. Images were captured on a Leica TCS SP8 confocal microscope and analyzed with Imaris software (Bitplane).

<u>*In vitro* cell culture</u>

Splenocytes and lymph node cells were isolated as described above, and naïve CD4 T cells were isolated using an EasySep Naïve CD4 T cell negative magnetic isolation kit (StemCell). $1x10^6$ cells were seeded into wells of 24 well plate coated with αCD3 and αCD28 antibodies for 2 days, in T cell culture media (T cell media without betamercaptoethanol and containing 10% FBS, 1x antioxidant supplement (Sigma Aldrich) and IL-2). Cells were then transferred to 6 well plate in 5mL media and were split 1:1 every 24 hours for the duration of culture. Sulfide supplement GYY4137 (Sigma Aldrich) was added to wells after transfer to 6 well plate. Untreated cells were given a vehicle control of spent GYY4137 at equal doses. Restimulation of cells for phospho-ERK staining was performed by coating cells in αCD3 Hamster IgG (Sigma Aldrich) and then cross linking with α-Hamster-IgG to initiate stimulation in 37°C water bath. Stimulation was halted with addition of 4% paraformaldehyde. Cells were then permeabilized with 100% ice cold methanol for 30 minutes before staining with intracellular phospho-ERK antibody (pT202/Y204 BD).

Proteomics

Naïve T cells were culture *in vitro* as described above, with a 3 day treatment with 100μM GYY4137. Cells were then subjected to the ProPerDP protocol to enrich for persulfidated proteins (*9*). Briefly, cells were lysed with RIPA buffer in a Bioruptor sonicator (Diagenode), then spun down to remove debris. The protein fraction was alkylated with 1mM EZ-link iodoacetyl biotin (Thermofisher), and desalted, with the total protein comprising fraction 1. Biotinylated protein was isolated with magnetic beads, fraction 2 containing the unbiotinylated fraction. The beads were treated with TCEP, and any released protein was the persulfidated fraction 3. The beads were then boiled in SDS-PAGE loading buffer to release fraction 4 represeting the thiol containing proteins without persulfidation. A Thermo Orbitrap Fusion Eclipse (Thermo Fisher Scientific, San Jose, USA) coupled to a Thermo UltiMate 3000 (Thermo Fisher Scientific) was used for LC-MS/MS experiments. 1 ug of total peptides were injected for LC-MS/MS analysis. Peptides were trapped on an Acclaim C18 PepMap 100 trap column (5 µm particles, 100 Å pores, 300 µm i.d. x 5 mm, Thermo Fisher Scientific) and separated on a PepMap RSLC C18 column (2 µm particles, 100 Å pores, 75 µm i.d. x 50 cm, Thermo Fisher Scientific) at 40 ºC. The LC steps were: 98% mobile phase A (0.1% v/v formic acid in H2O) and 2% mobile phase B (0.1% v/v formic acid in ACN) from 0 to 5 min, 2% to 35% linear gradient of mobile phase B from 5 to 155 min, 35% to 85% linear gradient of mobile phase B from 155 to 157 min, 85% mobile phase B from 157 to 170 min, 85% to 2% linear gradient of mobile phase B from 170 to at 172 min, 2% of mobile phase B from 172 to 190 min. Eluted peptides were ionized in positive ion polarity at a 2.1 kV spraying voltage. MS1 full scans were recorded in the range of m/z 375 to 1,500 with a resolution of 120,000 at 200 m/z using the Orbitrap mass analyzer. Automatic gain control and maximum injection time were set to standard and auto, respectively. Top 3sec data-dependent acquisition mode was used to maximize the number of MS2 spectra from each duty cycle. Higher-energy collision-induced dissociation (HCD) was used to fragment selected precursor ions with normalized collision energy of 27. MS2 scans were recorded using an automatic scan range with a resolution of 15,000 at 200 m/z using the Orbitrap mass analyzer. Data was analyzed with Proteome Discoverer (PD) normalized values, abundance ratio calculations, and statistical significance testing of protein abundances between conditions(sample/control). Total ion current (TIC) normalization (PD values) was used to correct technical variability, ensuring comparability across replicates. For statistical comparisons, PD applied two-sample t-tests (for pairwise conditions), calculating p-values and applying the Benjamini-Hochberg adjustment to control the false discovery rate, resulting in q-values/AdjP Values (adjusted P-values). When missing abundance data were seen in the label-free quantification data (LFQ), a criteria was set for at least two out of three values (for both controls and samples respectively) should be present before imputing the third missing value (in R through KNN imputation). To address this, we employed post-processing imputation steps outside PD to ensure more robust and interpretable statistical analyses.

Cholera toxin vaccination

Splenocytes from CD45.2 OT-II transgenic mice were isolated and total CD4 T cells were purified using magnetic separation (StemCell). $1x10^6$ T cells were injected retro-orbitally into CD45.1/45.2 mice. One day later, BSS treatment was started for 3 days. One day after BSS treatment ended, mice were gavaged with 10µg cholera toxin subunit B and 5 mg ovalbumin with 3% $NaHCO_3$ in $H_2O$. Alternately, mice were fed with SAA or defined AA diet for 2 weeks before initial vaccination, and kept on this diet for the duration of the experiment. Some SAA diet fed mice were also given NaHS containing drinking water as outlined above. Antigen specific response was assessed at 10 days post vaccination by ELISPOT, ELISA and flow cytometry for CD45.2+ OVA specific T cells.

ELISPOT and ELISA

Cells from ileum, mesenteric lymph node and spleen were isolated as outlined above and used for ELISPOT. 96-well plates (MultiScreen HTS, Millipore) were coated with 0.5 nM GM1-ganglioside overnight in PBS at 4°C. GM1-coated plates were further incubated with CTX (0.5 µg/ml) in PBS for at least 2 hours at room temperature. After washing the plates in PBS and blocking with 0.1% bovine serum albumin (BSA/PBS), isolated cells were seeded. The cells were incubated for 3 hours at 37°C and 5% $CO_2$. After thorough washing in PBS with 0.05% Tween 20, alkaline phosphatase (AP)–conjugated goat anti-mouse IgA or IgG (Southern Biotech) in 100 µl per well was added to the wells. The bound antibodies marking single antibody–producing cells were visualized by adding 50 µl per well of Sigmafast BCIP/NBT B5655 1 tablet per 10 ml deionized water (Sigma-Aldrich). ASCs were counted using an ImmunoSpot analyzer (CTL).

ELISAs were performed on isolated serum or small intestinal lumenal contents. Briefly, 96 well maxisorp ELISA plates (Nunc) were coated with CTX as described above, blocked with 0.1% BSA, and samples were serially diluted in 0.1% BSA/PBS and aliquots were added to corresponding subwells. The plates were kept at 4°C overnight, and after washing in PBS, the plates were incubated with AP–conjugated isotype-specific goat anti-mouse antibodies (Southern Biotechnology, Birmingham, AL). Plates were washed, and the AP substrate *p*-nitrophenyl phosphatase (Sigma-Aldrich), was added to each well. The reaction was read at 405 nm using a spectrophotometer.

Data Analysis and Statistics

Unless otherwise noted above, statistics were performed with Graphpad Prism 7. Flow cytometry statistics are presented as mean ± SEM or median with min to max using two-tailed unpaired t test for normally distributed data or Mann-whitney test for non-normal data. Normality of data sets was determined using Graphpad Prism. All experiments shown are representative of at least two independent experiments, with the exception of scRNAseq, ATACseq and metagenomics experiments.

#### Supplemental References


1. J. A. Hall *et al.*, Essential role for retinoic acid in the promotion of CD4(+) T cell effector responses via retinoic acid receptor alpha. *Immunity* **34**, 435-447 (2011).
2. E. Bolyen *et al.*, Reproducible, interactive, scalable and extensible microbiome data science using QIIME 2. *Nat Biotechnol* **37**, 852-857 (2019).
3. E. A. Franzosa *et al.*, Species-level functional profiling of metagenomes and metatranscriptomes. *Nat Methods* **15**, 962-968 (2018).
4. Y. Hao *et al.*, Dictionary learning for integrative, multimodal and scalable single-cell analysis. *Nat Biotechnol* **42**, 293-304 (2024).
5. A. M. Ackermann, Z. Wang, J. Schug, A. Naji, K. H. Kaestner, Integration of ATAC-seq and RNA-seq identifies human alpha cell and beta cell signature genes. *Mol Metab* **5**, 233-244 (2016).
6. A. M. Bolger, M. Lohse, B. Usadel, Trimmomatic: a flexible trimmer for Illumina sequence data. *Bioinformatics* **30**, 2114-2120 (2014).

7. A. Dobin *et al.*, STAR: ultrafast universal RNA-seq aligner. *Bioinformatics* **29**, 15-21 (2013).
8. S. Heinz *et al.*, Simple combinations of lineage-determining transcription factors prime cis-regulatory elements required for macrophage and B cell identities. *Mol Cell* **38**, 576-589 (2010).
9. E. Doka *et al.*, A novel persulfide detection method reveals protein persulfide- and polysulfide-reducing functions of thioredoxin and glutathione systems. *Sci Adv* **2**, e1500968 (2016).
10. X. Xie, P. Rigor, P. Baldi, MotifMap: a human genome-wide map of candidate regulatory motif sites. *Bioinformatics* **25**, 167-174 (2009).
