## Supplemental Figures for "Sulfide is a keystone metabolite for gut homeostasis and immunity"

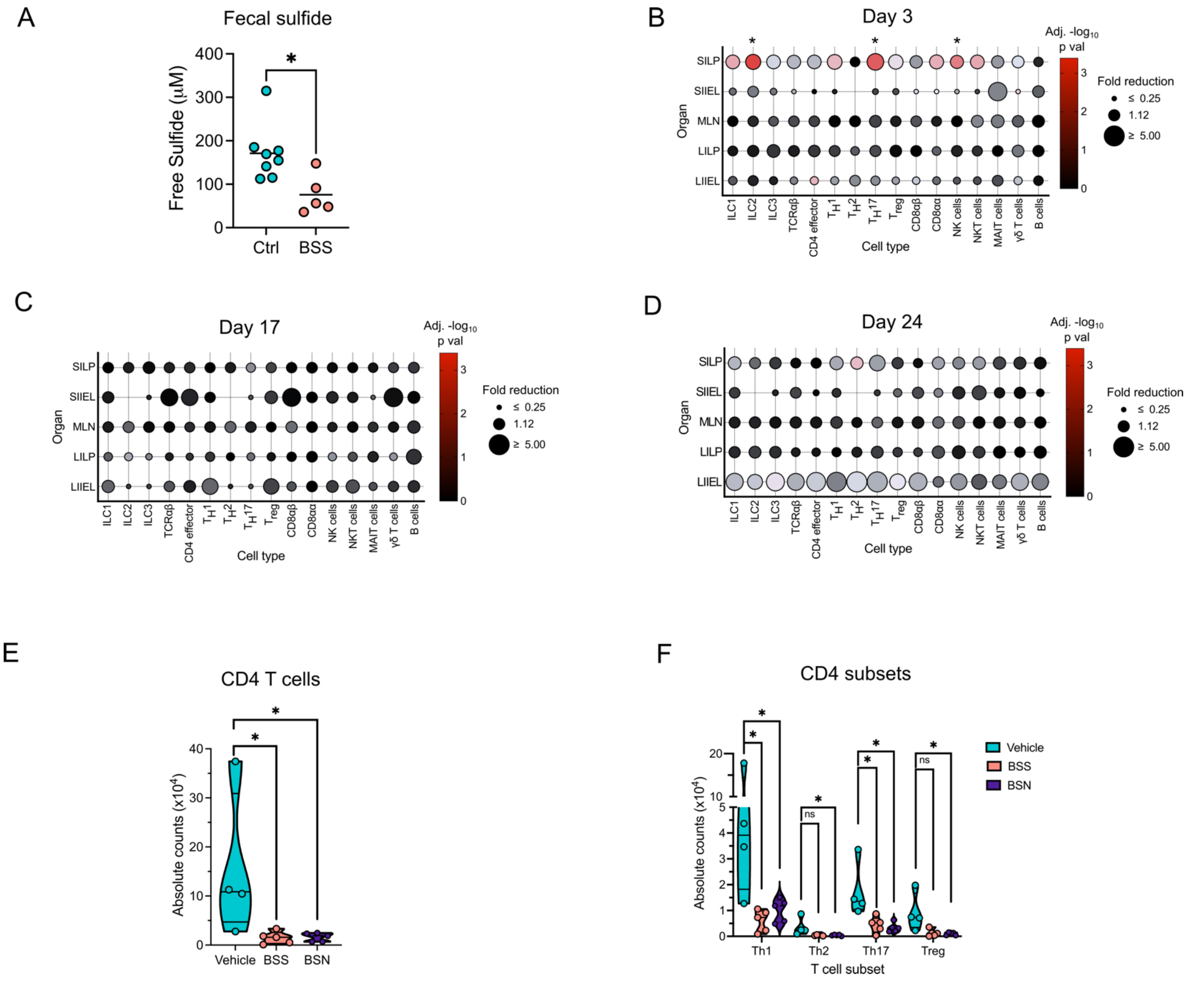


**Supplemental Figure 1.** (**A**) Free fecal sulfides in mice treated with control or BSS, 4 hours after final dose of BSS. (**B-D**), Kinetic of gut immune lymphocytes after BSS treatment**.** Cell counts for immune lymphocyte populations in small intestine lamina propria (SILP) and epithelium (SIIEL), mesenteric lymph nodes (MLN), large intestine lamina propria (LILP) and epithelium (LIIEL) at day 3 (B), 17 (C) and 24 (D) after start of BSS treatment. Size of circles indicates fold reduction of BSS treated mice compared to control and color indicates p value. (**E-F**), Small intestine CD4 T cell numbers and subsets after bismuth subnitrate treatment. Small intestine lamina propria CD4 T cell numbers (E) and CD4 T cell subsets (F) after BSS or bismuth subnitrate (BSN) treatment, at day 10 after start of treatment. * $p < 0.05$, ns not significant, two-tailed t-test, with Benjamini-Hochberg correction for B-D.

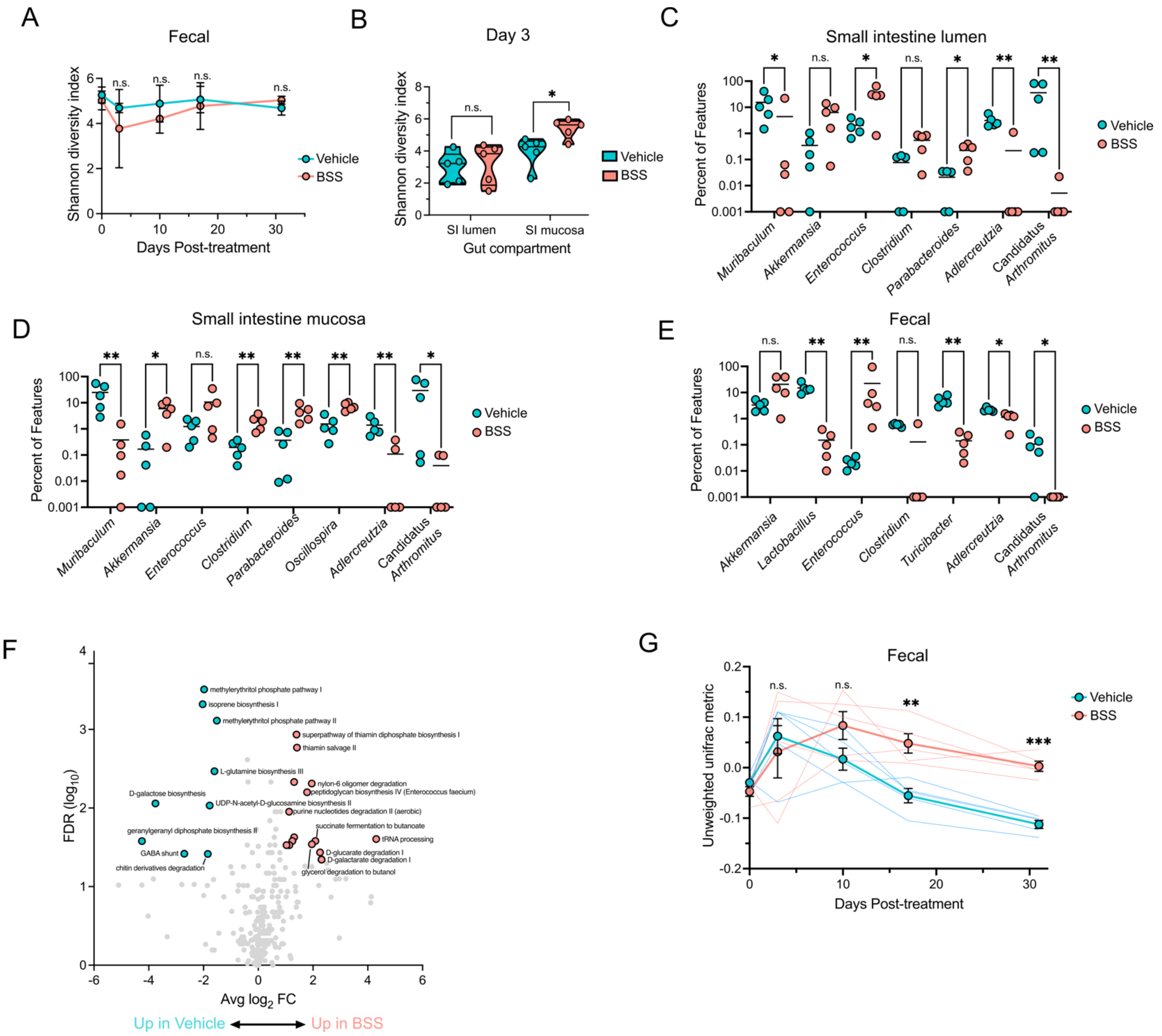


**Supplemental Figure 2. (A)** Alpha diversity of fecal microbiome over time after BSS treatment, as measured by Shannon diversity index after start of BSS treatment. (**B**) Alpha diversity of small intestine lumen and mucosa at day 3 after start of BSS treatment, as measured Shannon diversity index. **(C-E)** Identified genera in gut microbiome of BSS treated mice. Select genera identified by 16S rDNA sequencing of small intestine lumen (C) mucosa (D) or feces (E) microbiome in mice treated with BSS or vehicle at day 3 post treatment. **(F)** Metagenome alterations in BSS treated mouse microbiome. Fecal pellets collected from mice treated with BSS were submitted to shotgun metagenomics sequencing and analysis. Volcano plot of differentially expressed gene families, with significantly different genes highlighted in blue (increased in vehicle treated mice) or red (increased in BSS treated mice). (**G**) Beta diversity of fecal microbiome over time as measure by Axis 3 of PCA using the unweighted unifrac metric. ** $p < 0.01$, ** $p < 0.01$, *** $p < 0.001$, n.s. not significant, two-tailed t test (A-B), Mann-Whitney U test (C-F).

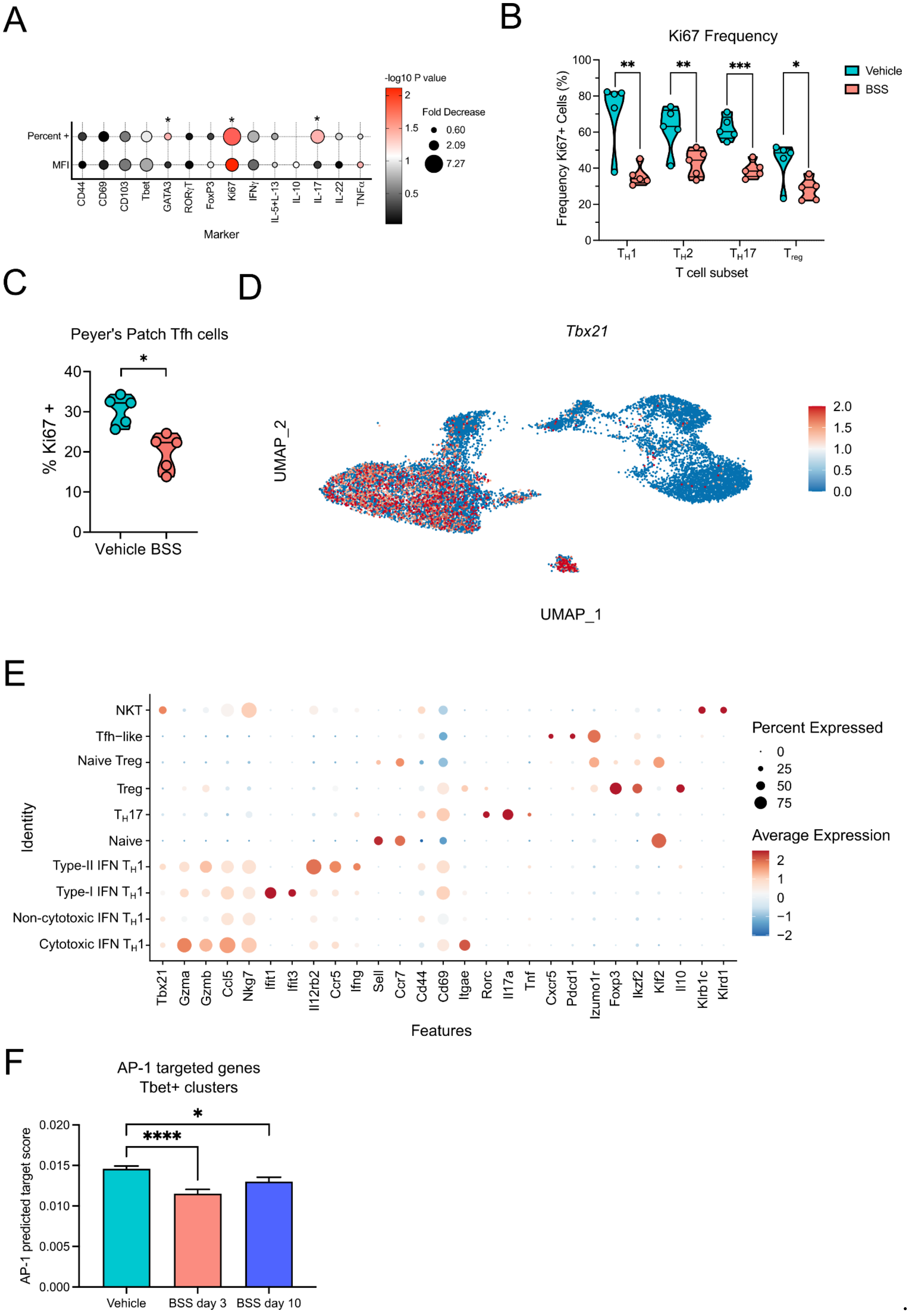

A
-log10 P value
Fold Decrease
0.60
2.09
7.27
Percent +
MFI
CD44
CD69
CD103
Tbet
GATA3
RORγT
FoxP3
Ki67
IFNγ
IL-5+L-13
IL-10
IL-17
IL-22
TNFα
Marker
B
Ki67 Frequency
Frequency Ki67+ Cells (%)
Vehicle
BSS
TH1
TH2
TH17
Treg
T cell subset
C
Peyer's Patch Tfh cells
% Ki67 +
Vehicle BSS
D
Tbx21
UMAP_2
UMAP_1
E
Identity
NKT
Tfh−like
Naive Treg
Treg
TH17
Naive
Type-II IFN TH1
Type-I IFN TH1
Non-cytotoxic IFN TH1
Cytotoxic IFN TH1
Tbx21
Gzma
Gzmb
Ccl5
Nkg7
Ifit1
Ifit3
Il12rb2
Ccr5
Ifng
Sell
Ccr7
Cd44
Cd69
Itgae
Rorc
Il17a
Tnf
Cxcr5
Pdcd1
Izumo1r
Foxp3
Ikzf2
Klf2
Il10
Klrb1c
Klrd1
Features
Percent Expressed
Average Expression
F
AP-1 targeted genes
Tbet+ clusters
AP-1 predicted target score
Vehicle
BSS day 3
BSS day 10

**Supplemental Figure 3. (A)** Analysis of CD4 T cell phenotypic markers at day 10 after start of BSS treatment, as represented by percent positive cells or median fluorescence intensity from flow cytometry. Size of circles indicates fold reduction of BSS treated mice compared to control and color indicates p value. **(B,C)** Percent Ki67 positive CD4 T cell subsets in small intestine lamina propria (B) or Peyer's Patches (C) at day 10 after start of BSS treatment. **(D)** UMAP of scRNAseq of ileal CD4 T cells, as in Figure 1J, with expression of *Tbx21* (Tbet) shown. (**E**) Dot plot of gene expression for each identified cluster in scRNAseq of ileal CD4 T cells. Genes shown were among the highest differentially expressed genes for each cluster and used for manual identification of each cluster. Size of circles indicates percent of cells expressing gene in each cluster, and color represents average expression for the gene in each cluster. (**F**) Module score of AP-1 activated genes in Tbet+ clusters. AP-1 score calculated based on 1576 predicted AP-1 target genes, from the MotifMap Predicted Transcription Factor Targets dataset (*10*). * $p < 0.05$, ** $p < 0.01$, *** $p < 0.001$, **** $p < 0.0001$, two-tailed t test with Benjamini-Hochberg correction (A), two-tailed t test (B,C), Mann-Whitney U test (F).

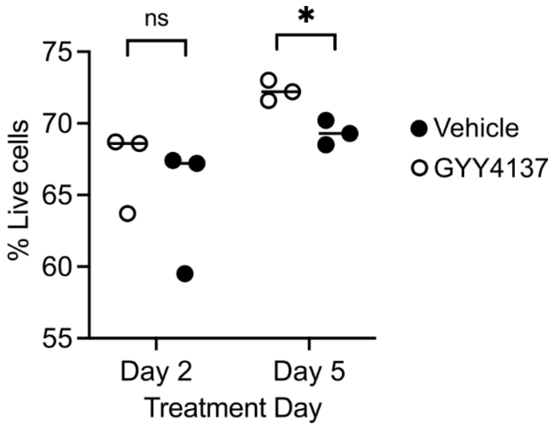


**Supplemental Figure 4.** Cell death measurement of CD4 T cells cultured with sulfide. CD4 T cells cultured with sulfide donor GYY4137 for indicated times, as in Figure 4 B-E, with amount of cell death indicated as measured using cell impermeable live/dead dye. * $p < 0.05$, ns not significant, two-tailed t test.

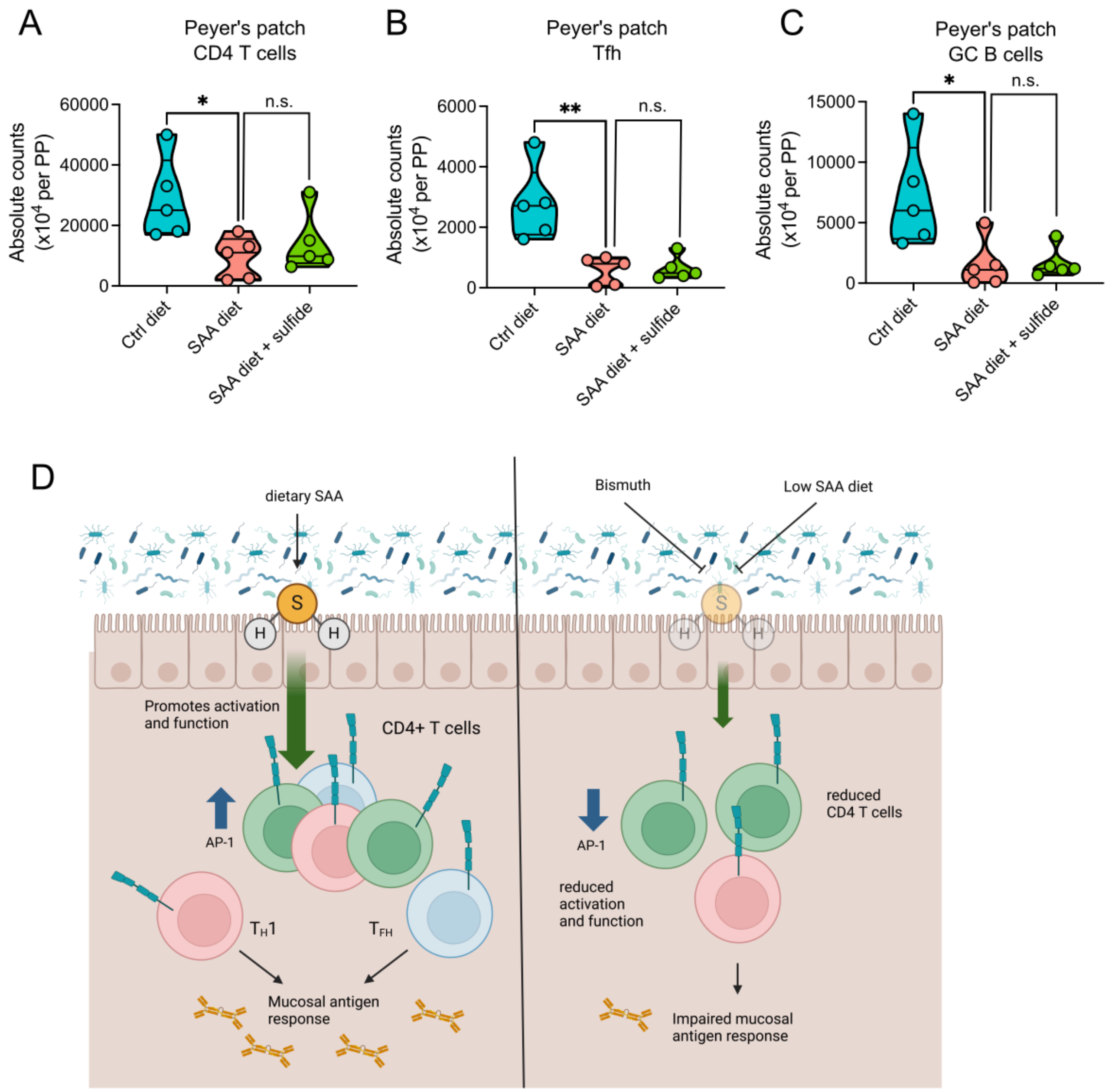


**Supplemental Figure 5. (A-C)** Mice treated with SAA diet +/- sulfide supplemented water and vaccinated, as in Figure 5I. Total Peyer's patch CD4 T cells (A), Tfh cells (B) or GC B cells (C) were quantified, represented as counts per Peyer's patch. (D) Schematic of the action of sulfide upon gut immune homeostasis and consequences of its disruption in the gut. * $p < 0.05$, ** $p < 0.01$, n.s. not significant, two-tailed t test.

### Supplemental Tables

**Table S1. Composition of diets.**

**Table S2. Significantly enriched proteins in proteomics of persulfidated protein fraction in CD4 T cells.**

**Table S3. Antibodies used in study.**
